# GenCore: Genomic distance estimation using Locally Consistent Parsing

**DOI:** 10.64898/2026.01.10.698768

**Authors:** Akmuhammet Ashyralyyev, Ege Sirvan, Salem Malikic, Tuğkan Batu, S. Cenk Sahinalp, Can Alkan

**Affiliations:** Dept of Computer Engineering, Bilkent University, Ankara 06800, Turkey; Cancer Data Science Laboratory, Center for Cancer Research, National Cancer Institute, National Institutes of Health, Bethesda, MD 20892, United States; Department of Mathematics, London School of Economics and Political Science, London, United Kingdom

**Keywords:** Locally consistent parsing, genome representation, comparative genomics

## Abstract

In the era of exponential data generation, a fast, consistent, and efficient string processing technique is necessary to represent extensive genomic data. One of the earliest string processing techniques, predating MinHash and minimizer-based sketching, is Locally Consistent Parsing (LCP). This technique partitions an input string and identifies short, exactly occurring substrings called *cores*, which collectively cover the input string while maintaining *Partition* and *Labeling Consistency*. The iterative application of LCP yields progressively longer cores in a compressed format, thereby substantially enhancing the efficiency of genomic sequence representation and subsequent downstream analysis.

We have previously developed Lcptools as the first iterative implementation of LCP for the DNA alphabet and demonstrated its effectiveness in identifying cores with minimal collisions. Here, we introduce GenCore, a computational method that leverages LCP cores for the first time to sketch and estimate genomic distances for closely related large genomes, and successfully reconstruct simulated progression trees. GenCore also successfully recapitulates primate phylogeny using both telomere-to-telomere (T2T) assemblies and the PacBio HiFi reads for assembly-free comparisons.

**Availability:** GenCore is available at https://github.com/BilkentCompGen/gencore

## 1. Introduction

Early studies in sequence-based comparative and evolutionary genomics date back to the 1970s, when algorithms for comparing strings were first introduced [1,2]. These foundational studies enabled the development of tools such as BLAST [3], which revolutionized sequence comparison by allowing rapid local alignments of genomic data. Classical phylogenetic tree construction methods, such as the Neighbor-Joining method [4], were employed to investigate evolutionary relationships.

As sequencing technologies became more affordable and more genomes were sequenced for evolutionary studies [5,6] and population studies [7,8], as well as for metagenomics [9], traditional alignment-based methods began to struggle with scalability due to the quadratic runtime complexity of sequence alignment algorithms. To address this challenge, alignment-free methods were introduced for genomic sequence distance estimation [10], which are further supported by *sketching techniques* to improve both execution time and memory requirements [11]. One of the most widely adopted sketching methods is MinHash [12], which is available in various implementations for sketch computation. The original approach utilizes multiple hash functions to determine the minimum hash value for each k-mer in the genomic dataset (i.e., the “document”) [12]. However, computing many hash functions can be computationally intensive when the input size is large. To overcome this, a simpler variant uses a single hash function and selects a subset of k-mers to create a compact signature of the sequence. The popular genome distance estimation tool `Mash` [13] utilizes this single-hash variant of MinHash, enabling resource-efficient phylogenetic analyses. Related sketching tools such as `Dashing` [14] and `Dashing 2` [15] instead use HyperLogLog- and SetSketch-based summaries to improve scalability and support efficient large-scale genome comparisons. However, traditional MinHash performs poorly on sets of highly dissimilar sizes. FracMinHash was later introduced as a recent modification to address this limitation, and implemented as `sourmash` [16].

Although `Mash`, `sourmash` and `Dashing 2` provide fast genome distance estimates, they struggle to accurately capture genuine relationships among very closely related genomes. This limitation is inherited from the fact that sketching methods [17], including MinHash [12] and FracMinHash [16,18], which *subsample* the set of available k-mers and therefore cannot fully capture the underlying sequence information. If sequence differences occur in regions not represented by selected k-mers, these variations will not be reflected in the distance calculations. Thus, `Mash` was optimized for **metagenomic data sets**^1^, which contain diverse and small genomes. To fully capture the distance between two **large and closely related genomes**, we need a method that produces sketches with ungapped coverage across consecutive substrings, so that sequence variation is represented throughout the genome rather than only in sampled regions.

Here, we propose leveraging Locally Consistent Parsing (LCP) [20,21] to parse genomic sequences into substrings called *cores*, and assign them labels to build a *comprehensive* representation set. Unlike subsampling-based sketching methods, LCP-based parsing preserves coverage of the underlying sequence by representing all regions through parsed substrings. This property makes it well-suited for detecting localized differences that may otherwise be missed by sparse *k*-mer sketches.

In this work, we use an iterative LCP-based approach to partition a string into progressively higher-level, fewer substrings while maintaining a representation of the full sequence. This hierarchical representation allows sequence changes at different scales to contribute to the distance computation. We first introduced this LCP-based string partitioning strategy in LCPan [22], where we used LCP for variation graph construction. Here, we present an alternative use case for LCP partitioning and adapt it to address limitations of sketch-based distance estimation in phylogenetic inference.

## 2. Methods

### 2.1 Locally Consistent Parsing

We recently developed a toolkit to process an input string named Lcptools [22] based on the Locally Consistent Parsing (LCP) technique [20,21]. LCP method partitions an input string into similarly-sized substrings called *cores*, with two guarantees: 1) *partition consistency*, i.e., different parts of the input string with the same content are partitioned the same way, and 2) *labeling consistency*, i.e., cores that represent the same underlying sequence are assigned the same label. Furthermore, LCP can be applied iteratively across *levels*, where the cores at each level encode increasingly longer substrings. As the hierarchy progresses, short, abundant lower-level cores are merged into fewer, longer higher-level cores, reducing the number of cores by a constant factor of approximately *c* ∼ 2.34 between consecutive levels [22].

Briefly, Lcptools detects *core substrings* using the following criteria. First, a length-3 substring *w* = *xyz* is classified as an **LMIN core** when its middle character *y* is a *proper* local minimum. That is, assuming *x* ≠ *y* and *y* ≠ *z*, the inequalities *x > y* and *y < z* must both be satisfied. Second, a length-3 substring *w* = *xyz* is classified as an **LMAX core** when its middle character *y* is a proper local maximum, while neither neighboring character *x* nor *z* is a local minimum. More specifically, if *w* = *xyz* occurs inside the larger context *sxyzt*, then *y* must satisfy *x < y* and *y > z*, together with the additional conditions *s* ≤ *x* and *z* ≥ *t*. Third, a substring *w* with |*w*|*>* 3 is classified as a **RINT core** when all characters except its first and last are the same. Formally, for *w* = *xy*^*i*^*z*, where *i >* 1,*x* ≠ *y*, and *y* ≠ *z*, the substring satisfies the RINT core condition. Fourth, a substring *w* = *za*_1_ … *a*_*n*_*k* is classified as an **SSEQ core** when it connects two already identified cores, with both of its endpoints *z* and *k* serving as core boundaries. This endpoint condition is equivalent to requiring the sequence between the two cores to be strictly monotonic; that is, either *z < a*_1_ *<* … *< a*_*n*_ *< k* or *z > a*_1_ *>* … *> a*_*n*_ *> k*, where *n*≥1. Therefore, an SSEQ core represents a stranded sequence whose two ends are cores and whose characters follow a strictly increasing or strictly decreasing order.

Each core is represented using the 2-bit encoding of its corresponding substring (i.e., A=00, C=01, G=10, and T=11). However, applying multiple rounds of LCP rules directly can lead to excessively long bit strings. To address this, Lcptools applies Deterministic Coin Tossing (DCT) [23] as a compression step before each further LCP iteration. For every core, DCT locates the first bit position at which the core differs from its immediate left neighbor and appends that differing bit. This produces a compact, comparable alphabet that allows LCP to be applied repeatedly while maintaining the required correctness guarantees. Together, these rules partition the string into nearly balanced segments and ensure that identical substrings receive consistent representations wherever they occur. Because LCP runs in linear time and requires linear space, it is well-suited for large-scale genomic data processing.

### 2.2 Genomic distance estimation with Gen Core

GenCore, an alignment-free method, leverages LCP to quickly estimate genomic distances between multiple species and/or tumor cell lines. GenCore follows a strategy similar to `Mash` [13], which uses the MinHash [12] method to calculate the Jaccard similarity of the k-mer compositions of two data sets. However, `Mash` only extracts a subset of sketches (1000 by default) with minimal values from the calculated hash function. Thus, the pairwise comparison between those sketches yields an estimated Jaccard similarity score, which is then used to calculate the *Mash distance*, which correlates with average nucleotide identity (ANI). GenCore supports input from both assembled genomes and unassembled read data sets generated by PacBio HiFi.

In contrast, GenCore estimates genomic distances using the framework outlined in Algorithm 1. Briefly, GenCore processes each genomic data set up to a specified LCP level and stores the labels of its partitions as a sketch. It then calculates the Jaccard distances and the *p*-distances for each pair of sketches to construct the distance matrix.

#### Algorithm 1

Genomic distance estimation with GenCore

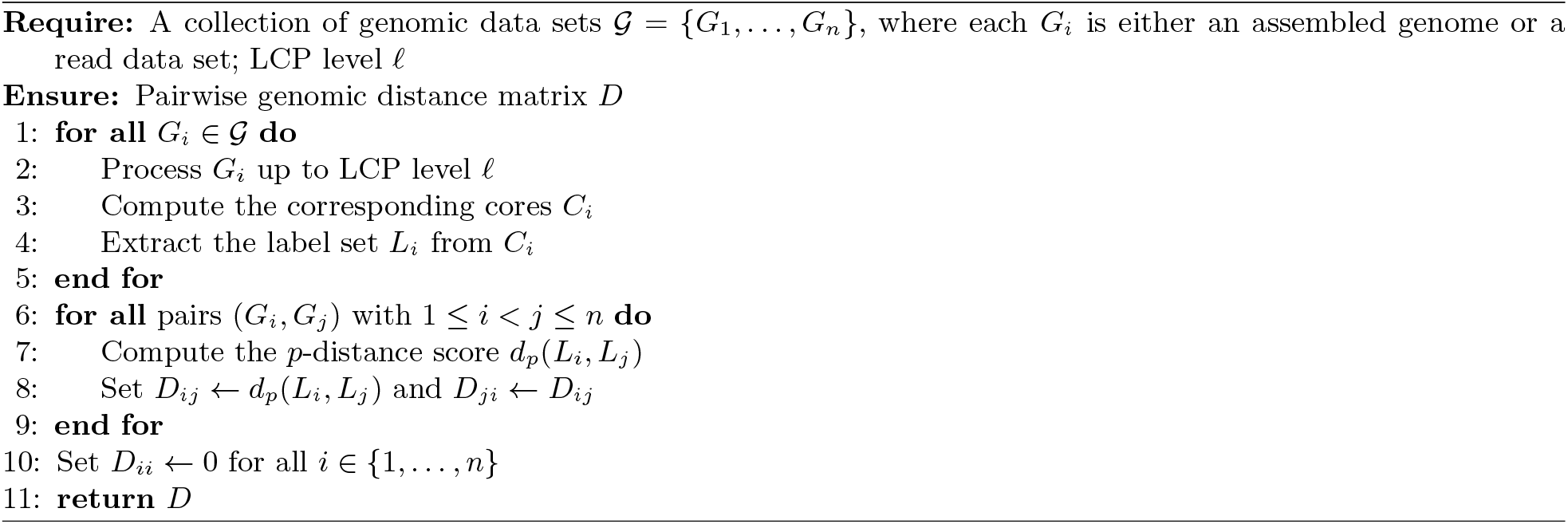

If multiple genomic data sets are provided for a phylogenetic analysis, GenCore extends the distance calculation for all pairs, and then constructs a phylogenetic tree using the Neighbor-Joining method. The core idea behind LCP cores is to accurately represent the underlying sequences. Errors can lead to different partitions and sub-cores, which in turn cause variations in the core labels, ultimately resulting in different upper-level core labels. Therefore, the LCP method is particularly well-suited for low-error reads, such as those generated by PacBio HiFi.

Similar to `Mash`,GenCore first calculates the Jaccard similarity. Note that for two sets

*A* and *B*, the Jaccard similarity is defined as follows:

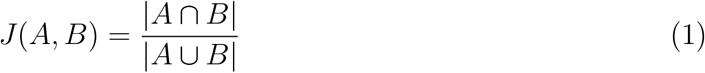

Assuming the simple evolutionary model under a Poisson model results in a *p*-distance (otherwise known as Mash-distance), which can be approximated using the Jaccard score [24] that we refer to simply as *J* :

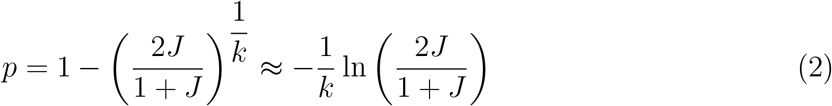

In our model, each LCP core label is treated analogously to a *k*-mer for compression and distance-estimation purposes. Thus, Equation 2 follows the standard *p*-distance formulation used for *k*-mer-based comparisons, with the *k*-mer length term replaced by the corresponding core-length estimate. Note that *k* is a fixed k-mer size, as used by `Mash` [13]. However, the lengths of the cores generated by LCP vary. To address this, we use the average core length at the applicable LCP level instead. Finally, GenCore constructs the phylogenetic trees from the distance matrix using Biopython [25].

Theorem 1 below provides a theoretical analysis that supports the observed accuracy of GenCore in genomic distance estimation. The theorem is stated in terms of base substitutions, but an analogous statement can be made for more general random mutations, including insertions and deletions.

#### Theorem 1

*Let S be a genomic sequence of length n that is generated uniformly at random. Let sequence S*^*′*^ *be obtained by applying base substitutions to S at a rate of ε: each position in S is chosen independently at random with probability ε and the base is replaced with a base that is chosen uniformly at random (including the same base currently at that location). Let ℓ be the level of the LCP application that satisfies that the average core length k is at most* (*Cε*)^*−*1^ *for some constant C >* 1. *The expected p-distance between S and S*^*′*^ *calculated using level-ℓ cores is ε, where the expectation is calculated over the generation of S and the substitutions*.

*Proof*. Let *X* be the number of level-*ℓ* cores that are changed by a single random substitution as described above. We note that E[*X*] = *O*(1). As a result, for sets *A* and *B*, respectively, of level-*ℓ* cores obtained from *S* and *S*^*′*^, respectively, the Jaccard similarity between *A* and *B* is

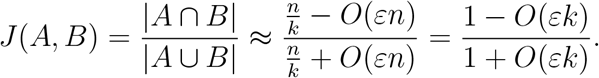

Next, plugging this into the *p*-distance definition in Equation 2, we get

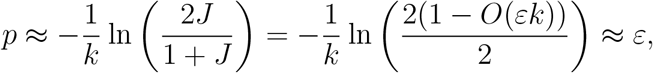

where the last approximation follows from *k* ≤ (*Cε*)^*−*1^ and that, for small *x*, ln(1 − *x*) ≈−*x*.

We now turn to analyze E[*X*]. Firstly, we note that a base substitution can change at most three consecutive cores at level 1. In turn, these changes to the cores affect partitioning into level-2 cores, where up to 7 level-1 cores are involved. As a result, at most 3 level-2 cores are affected by this base substitution. Repeating this argument for each layer, we establish that the number of cores changed at each level remains constant. Furthermore, by keeping the average core length small enough (*k* = *O*(*ε*^*−*1^)), we ensure that separate substitutions do not interact with each other, so that most of the substitutions are observed as separate level-*ℓ* cores that are changed.

### 2.3 Assembly-free phylogenetic analyses with GenCore

GenCore supports phylogenetic analysis directly from unaligned sequencing reads generated by the PacBio HiFi platform, bypassing the need to build genome assemblies and perform whole-genome alignments, or map reads to a reference.

To enable this assembly-free capability, we extend the original methodology of GenCore with two key enhancements: (1) processing both original and reverse-complements of the reads to ensure strand-oblivious information representation, and (2) removing LCP cores with frequencies less than the average read depth of the sequencing data to reduce noise introduced by sequencing errors. These steps collectively preserve the full informational content of the reads while improving robustness against sequencing artifacts.

## 3 Results

We tested GenCore using both simulated and biological sequence data. We estimated genomic distances of tumor genomes and then built phylogenetic trees to compare with the underlying simulation. We also analyzed primate genomes, both using T2T assemblies and using unassembled PacBio HiFi read sets. In all our experiments, we compared our results with those from `Mash`, `sourmash` [26], and `Dashing 2`.

### 3.1 Tumor phylogeny inference using GenCore

We first evaluated GenCore on a tumor phylogeny simulation, comparing the inferred phylogenetic tree with the simulated ground truth tree. For this purpose, we used a Monte Carlo-based phylogeny simulator (`mksamples`) [27] to generate a tree with 20 leaf nodes. We then used a phylogeny-guided tumor evolution simulator (PSiTE) [28] to generate sub-clonal genomes that “evolved” following the simulated phylogeny. The simulated genomes were generated from the human reference genome (GRCh38) using phased mutations in the HG001 (NA12878) genome [29]. We did not include the Y chromosome in the simulations, as HG001 is a female sample. Each subclonal tumor genome included single-nucleotide variations (SNVs) and copy number variations (CNVs, i.e., deletions). The average length of CNVs was 2 Mbp, whereas the largest CNVs were 4 Mbp.

We used GenCore to estimate pairwise genomic distances based on the Poisson model for random-site mutations across the 20 simulated tumor genomes, and we repeated the experiment for LCP levels 4 to 8. Processing all simulated diploid genomes with Lcptools to extract core labels took an average of 14 minutes, using 4 CPU threads. Pairwise comparisons for all 20 genomes were completed in a few seconds, substantially faster than performing whole-genome alignments. We also used `Mash` [13], a popular alignment-free method that predicts genomic distances using MinHash sketching [12], to compare with the GenCore results across various sketch sizes. Estimating genomic distances for all pairs of 20 tumor genomes using `Mash` took 14 minutes, using 4 CPU threads with a 5-million sketch size. However, we reiterate that `Mash` was primarily designed for metagenome distance estimation, where GenCore is optimized for comparing closely related large genomes.

We then constructed phylogenetic trees using the Neighbor-Joining module of the Biopython package [25] using the predicted genomic distance matrices. (Figure 1).

**Fig. 1.**
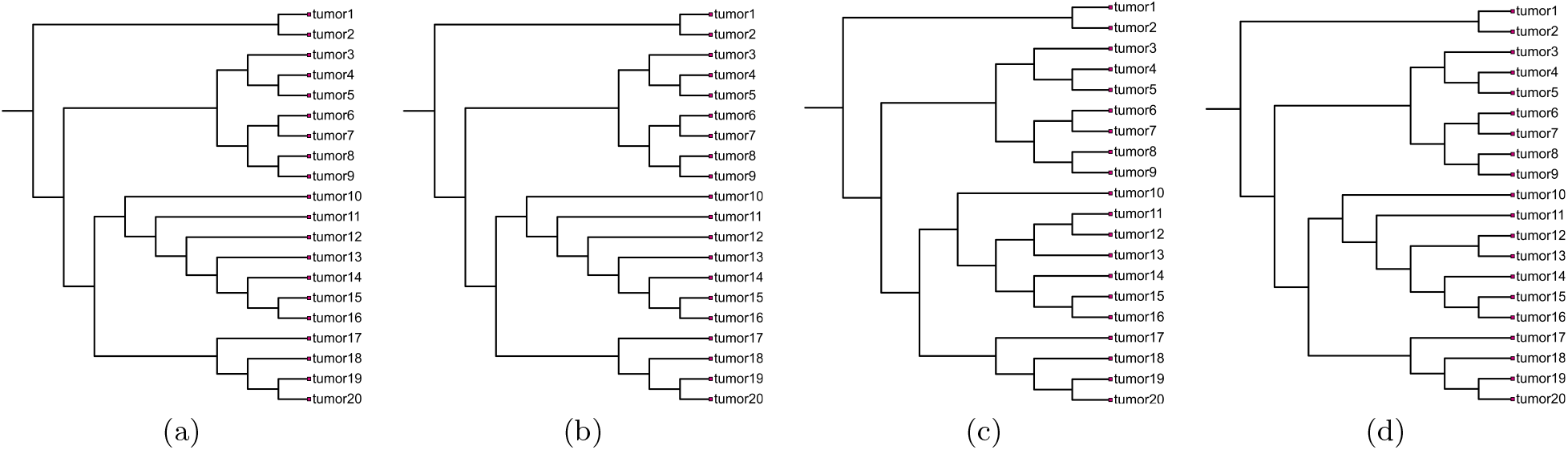
Phylogenetic trees constructed from simulated tumor genomes using Gen Core compared with Mash. (a) Simulated tree (ground truth), (b) Gen Core tree at LCP level 7, (c) tree generated by `Mash` with a sketch size of 5 million, and (d) tree generated by `Dashing 2` with a sketch size of 5 million. The phylogenetic tree estimated by Gen Core is isomorphic to the simulated tree. Additional Gen Core trees generated at different LCP levels, as well as trees generated by `Mash` and `Dashing 2` using different sketch sizes, are provided in Supplementary Figures fig. S1, fig. S2, and fig. S3.

We observed that GenCore generated trees that are isomorphic to the true tree (Figure 1a) in LCP level 6 (Figure 1b). GenCore accurately reconstructed the simulated tumor phylogeny across all evaluated LCP levels (4-8), producing trees that were isomorphic to the simulated ground-truth topology (Figure 1a,b). In contrast, `Mash` and `Dashing 2` failed to recover the correct phylogeny using its default sketch sizes, estimating all simulated tumor cells to be effectively identical. Increasing the sketch size progressively improved the inferred topology but did not recover the ground-truth tree, even with a sketch size of 5 million (Figure 1c,d). We also evaluated `sourmash`, which reported Jaccard similarities of 100% for all genome pairs and therefore failed to distinguish among the simulated tumor genomes. Consequently, we excluded the `sourmash` results from further analysis of this data set.

To quantitatively compare the predicted trees with the ground truth, we used the Robinson– Foulds (RF) [30] distance. The RF distance between two trees is defined as the number of non-trivial splits present in one tree but not the other, i.e., *d*_RF_(*T*_1_,*T*_2_)= |*S*(*T*_1_) △*S*(*T*_2_) | . Further, the normalized RF distance (nRF) is calculated by dividing the RF distance by the maximum possible RF distance for the number of fully resolved unrooted binary trees with *n* leaves, i.e., 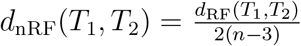 Table 1 summarizes the RF and nRF distances of the predicted trees with the ground truth, resource usage, and run times for the tumor phylogeny inference experiments.

**Table 1.**
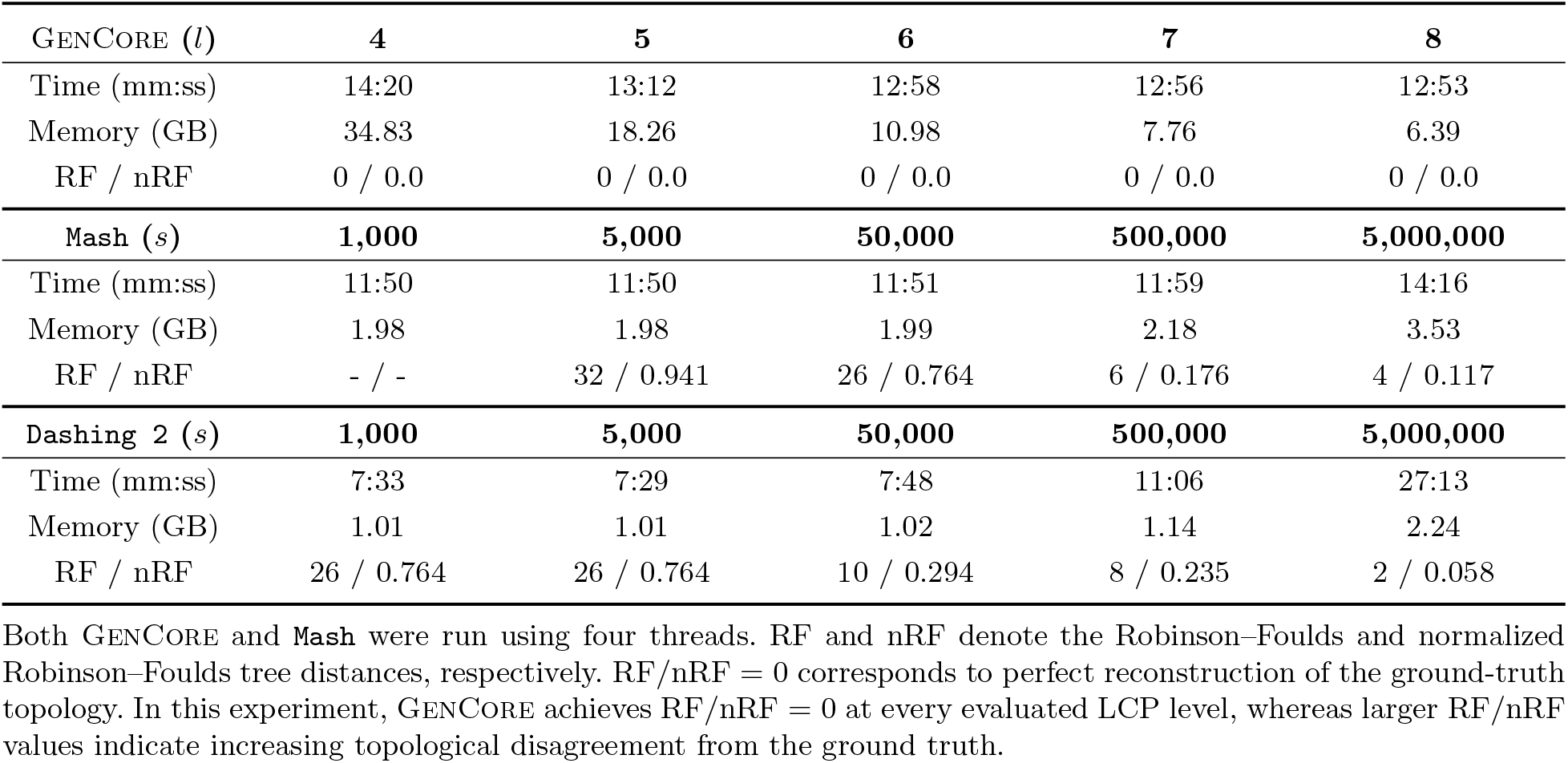
Resource usage, execution time, and phylogenetic accuracy for simulated tumor cells using GenCore, Mash and Dashing 2.

| GENCORE (*l*) | 4 | 5 | 6 | 7 | 8 |
| --- | --- | --- | --- | --- | --- |
| Time (mm:ss) | 14:20 | 13:12 | 12:58 | 12:56 | 12:53 |
| Memory (GB) | 34.83 | 18.26 | 10.98 | 7.76 | 6.39 |
| RF / nRF | 0 / 0.0 | 0 / 0.0 | 0 / 0.0 | 0 / 0.0 | 0 / 0.0 |
| Mash (*s*) | 1,000 | 5,000 | 50,000 | 500,000 | 5,000,000 |
| Time (mm:ss) | 11:50 | 11:50 | 11:51 | 11:59 | 14:16 |
| Memory (GB) | 1.98 | 1.98 | 1.99 | 2.18 | 3.53 |
| RF / nRF | - / - | 32 / 0.941 | 26 / 0.764 | 6 / 0.176 | 4 / 0.117 |
| Dashing 2 (*s*) | 1,000 | 5,000 | 50,000 | 500,000 | 5,000,000 |
| Time (mm:ss) | 7:33 | 7:29 | 7:48 | 11:06 | 27:13 |
| Memory (GB) | 1.01 | 1.01 | 1.02 | 1.14 | 2.24 |
| RF / nRF | 26 / 0.764 | 26 / 0.764 | 10 / 0.294 | 8 / 0.235 | 2 / 0.058 |

Both GenCore and `Mash` were run using four threads. RF and nRF denote the Robinson–Foulds and normalized Robinson–Foulds tree distances, respectively. RF/nRF = 0 corresponds to perfect reconstruction of the ground-truth topology. In this experiment, GenCore achieves RF/nRF = 0 at every evaluated LCP level, whereas larger RF/nRF values indicate increasing topological disagreement from the ground truth.

#### Accuracy of GenCore -estimated distances

We evaluated the correlation between sequence distances and the *p*-distances estimated by GenCore using cores at varying LCP levels. Because tumor genomes are simulated using the human genome as a baseline, retaining its substantial size and complexity (i.e., repeats and segmental duplications), directly calculating the edit distance is not trivial. Consequently, we employed the mutational divergence^2^ between pairwise tumor sublines as a proxy for Average Nucleotide Identity (ANI)[31]. Given that the simulated genome sizes remain approximately uniform, the count of distinct mutations serves as a reliable correlate for ANI. We computed the mutational differences for each tumor genome pair and analyzed their correlation with p-distances derived by GenCore using LCP core levels 4 through 9 (Supplementary Figure fig. S4). We also conducted a parallel assessment using Jaccard distances (Supplementary Figure fig. S5). In all instances, the comparisons showed a strong correlation (*R*^2^ ≥ 0.9) after LCP level 4.

### 3.2 Primate phylogeny using GenCore

Next, we used GenCore on biological data to build the phylogenetic tree of primates. We downloaded six finished diploid non-human primate genomes and a diploid human genome (HG002) generated by the T2T Consortium [6]. We used GenCore to estimate genomic distances and build phylogenetic trees, and repeated our experiment for LCP levels 4 to 6 (Figures 2 and S6). LCP cores were extracted for all seven diploid primate genomes in ∼5 minutes with the LCP level 4, and ∼4 minutes with the LCP levels 5 and 6 using seven threads.

**Fig. 2.**
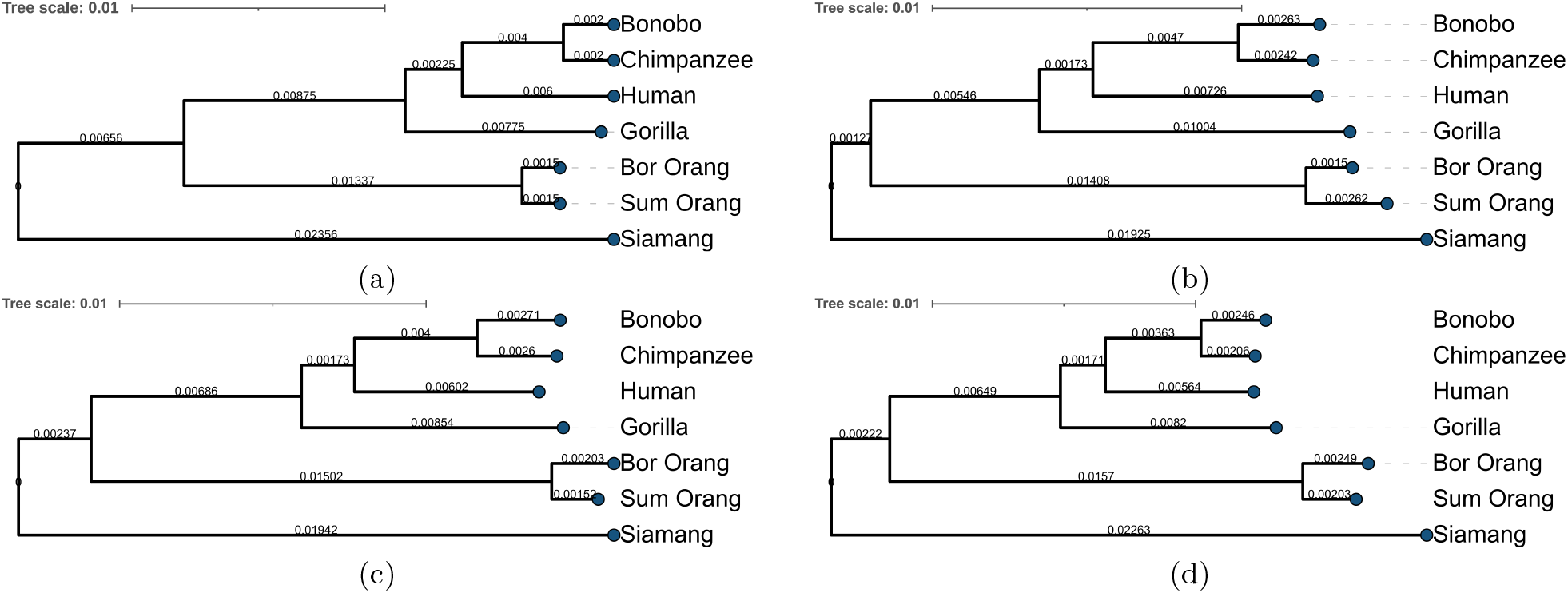
Phylogenetic trees constructed using complete primate genomes. (a) known phylogeny, (b) tree constructed using GenCore at LCP level 6, (c) tree constructed using Mash with sketch size 1000, (d) tree constructed using Sourmash with Jaccard similarity scores converted into Mash distances with k-mer size 21.

We observed that GenCore -generated trees are isomorphic to the known phylogeny, although the estimated branch lengths slightly differed among different settings. Note that `sourmash` computes only the Jaccard similarity scores. Hence, for each score, we applied Equation 2 to convert Jaccard similarity scores into p-distance estimates and computed a phylogenetic tree from the resulting values.

We analyzed branch lengths to better understand the differences among the phylogenies generated by GenCore at different LCP levels. In the absence of a whole-genome alignment-based tree, we used the evolutionary distances among the species to calculate the branch lengths in the Neighbor-Joining tree for the known phylogeny [6,32]. As we increased the LCP level, the distances estimated by GenCore converged to the known evolutionary distances at LCP level 6, as shown in Figure 2. We annotated the estimated distances in the branches and provided all trees in the Supplementary Figure fig. S6.

### 3.3 Assembly-free phylogeny inference using GenCore

Lastly, we inferred a phylogenetic tree using only raw HiFi reads from primate genomes, excluding the human genome, based solely on the sequencing data from Yoo et al. [6]. By default, we set the minimum LCP core-frequency threshold to 32, corresponding to the lowest diploid depth of coverage among the datasets, to reduce noise from sequencing errors (Supplementary Table table S1). To assess the effect of this filtering threshold, we repeated the analysis at LCP level 6 across thresholds ranging from no filtering to 200 and present our findings in Figure 4. The weighted Robinson–Foulds distance to the known phylogeny was minimized at a threshold of 16, which corresponds to half of the minimum diploid coverage. This is consistent with the expectation that, under approximately even read distribution across DNA strands, GenCore can retain LCP cores supported by both strands while removing low-frequency error-derived signals. Beyond a threshold of approximately 30, the weighted Robinson–Foulds distance began to increase, indicating that overly stringent filtering removes phylogenetically informative signal and eventually disrupts the inferred tree topology.

We employed LCP levels 5 and 6, as illustrated in Figure 3a and Figure 3b, which previously yielded highly accurate results in our analyses. We observed improved memory and runtime performance with higher LCP levels. Detailed benchmarking analyses are presented in Table 2. In total, 1.2 TB of sequencing data were processed. Data size properties are provided in Supplementary Table table S1.

**Table 2.**
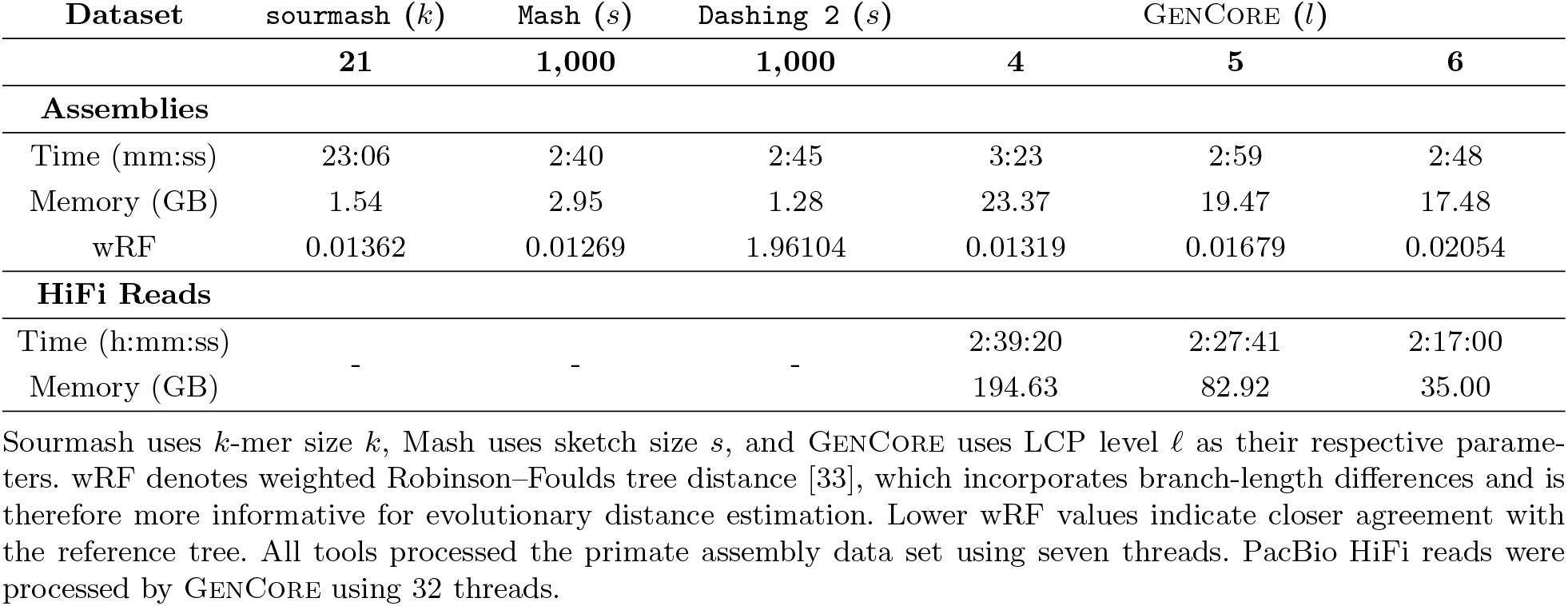
Resource usage, execution time, and phylogenetic distance metrics for computing phylogeny of the T2T primate genomes.

**Fig. 3.**
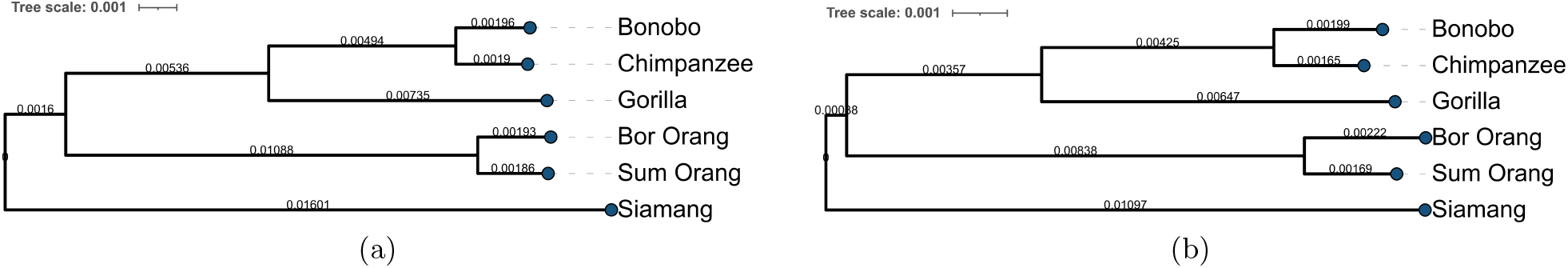
Phylogenetic trees constructed using PacBio HiFi reads generated from primate genomes. Trees are constructed with (a) LCP level 5, (b) LCP level 6. All LCP levels generated the true phylogeny, while the weighted RF distance increased at higher LCP levels.

**Fig. 4.**
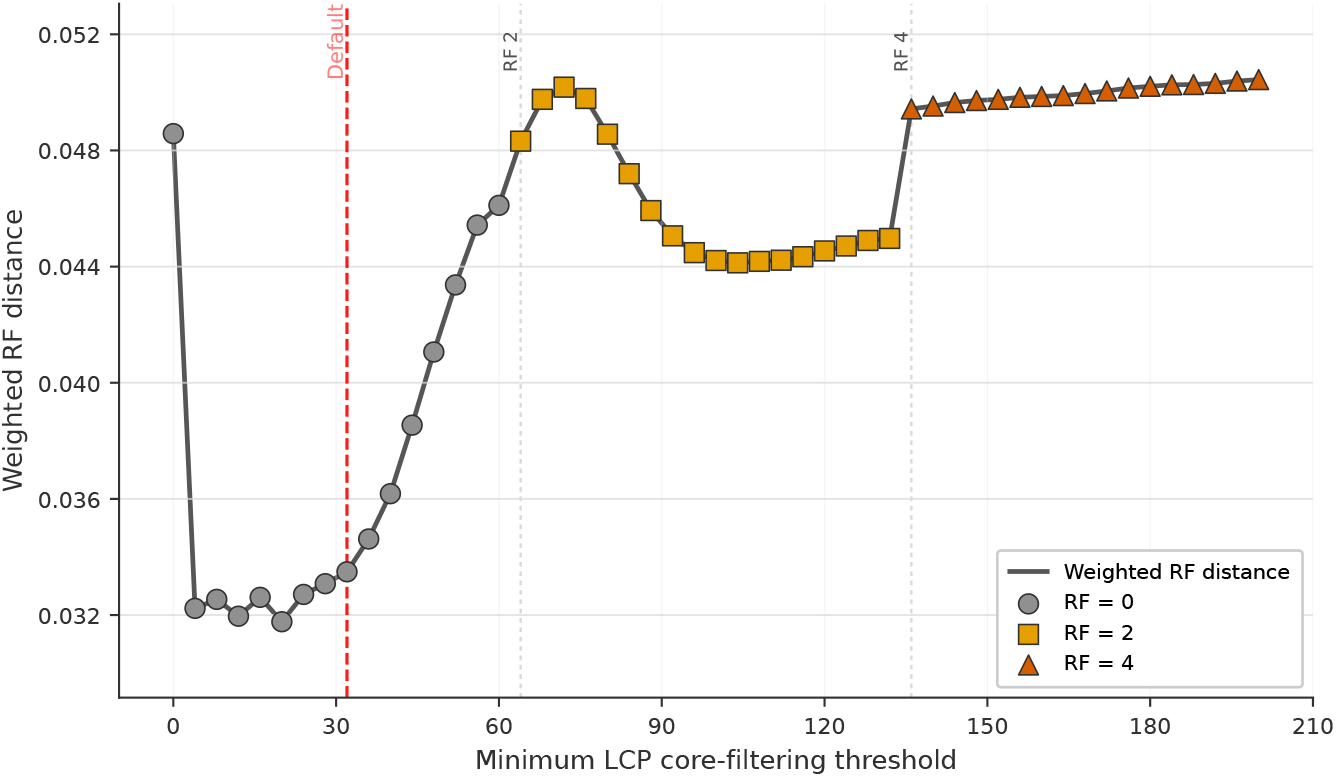
Effect of frequency-based LCP core filtering threshold on primate phylogeny accuracy.

## 4 Discussion

In this article, we presented GenCore, a method for representing large genomes in an ungapped and efficient manner using locally consistent parsing (LCP). Our results demonstrate that LCP-based genome representations can accurately capture evolutionary relationships while allowing users to control the granularity of comparisons via LCP levels. In particular, higher LCP levels substantially reduce memory consumption while still producing isomorphic phylogenetic trees for large eukaryotic genomes in our tumor simulations. Moreover, GenCore was successful not only in reconstructing phylogenetic trees but also in capturing true evolutionary distances among primate genomes. These results suggest that tumor genome relationships can be analyzed assembly-free, reducing the need for complete genome assembly before downstream evolutionary analysis.

In our experimental analyses, we observed a distinct relationship between LCP level selection and the genomic distance of the samples being processed. Specifically, for closely related genomes (intra-species), the optimal LCP level for accurately reconstructing the true phylogeny was found to be 7–8. On the other hand, when analyzing more distantly related genomes, such as those from different species, selecting LCP levels of 5–6 was more effective for capturing the genomic diversity. LCP levels exceeding these optimal ranges tended to yield deteriorated results. For assembly-free analyses, the exclusive use of PacBio HiFi reads, with low error rates, enabled the accurate resolution of true intra-species phylogenetic relationships. We leave a formal theoretical analysis of the relationship between LCP level, sequence divergence, and phylogenetic accuracy for future work.

Although GenCore is slower than and requires more memory than `Mash` and `sourmash`, it provides superior accuracy in estimating the genomic distances of closely related large genomes. The performance overhead is primarily due to LCP processing, which generates a large number of LCP cores at lower levels, each storing associated metadata. To mitigate this, a divide-and-conquer approach could be employed to partition the workload, potentially reducing both execution time and memory usage. Additionally, `Mash` and `sourmash` subsample from k-mers, which contribute to their speed and low memory footprint, while performing *lossy* comparisons. On the other hand, GenCore effectively compares entire genomes at user-defined coarseness (i.e., LCP levels), which improves accuracy but incurs runtime and memory costs. Note that at higher LCP levels, GenCore requires less memory and runs faster due to reduced data movement and fewer core comparisons.

In the context of raw read analyses, we believe that a more efficient filtering mechanism that does not require all data to be retained until the final step could further improve memory efficiency. Additionally, the filtering frequency thresholds are selected based on pre-calculated depth-of-coverage. Future work will focus on extending GenCore to dynamically determine appropriate frequency thresholds during processing, making the tool more adaptive and less reliant on prior depth information. Another future direction will be extending GenCore for metagenomic classification.

## 6 Author contributions statement

A.A., S.C.S., S.M., and C.A. developed the methods and conceived the experiments. E.S., and T.B. developed the theoretical framework for the relationship between LCP core levels and p-distance estimation. A.A. implemented the methods and conducted the experiments. A.A., S.C.S., and C.A. wrote and reviewed the manuscript.

## Acknowledgments

We thank all Alkan Lab members for their insightful comments and suggestions. Special thanks to Ricardo Román-Brenes for technical support during implementation. We also thank the members of the SAFARI Group, especially Nika Mansouri Ghiasi and Konstantina Koliogeorgi, for valuable discussions and exchange of ideas. This work was partially supported by the European Union’s Horizon Programme for Research and Innovation under grant agreement No. 101047160, project BioPIM, and the Intramural Program of the National Cancer Institute.

## Footnotes

1 Though, it performs well with large primate genomes with high sequence divergence (*>*1.5%) [19].

2 As simulated using PSiTE [28].

