## Supplementary Material for "GenCore: Genomic distance estimation using Locally Consistent Parsing"

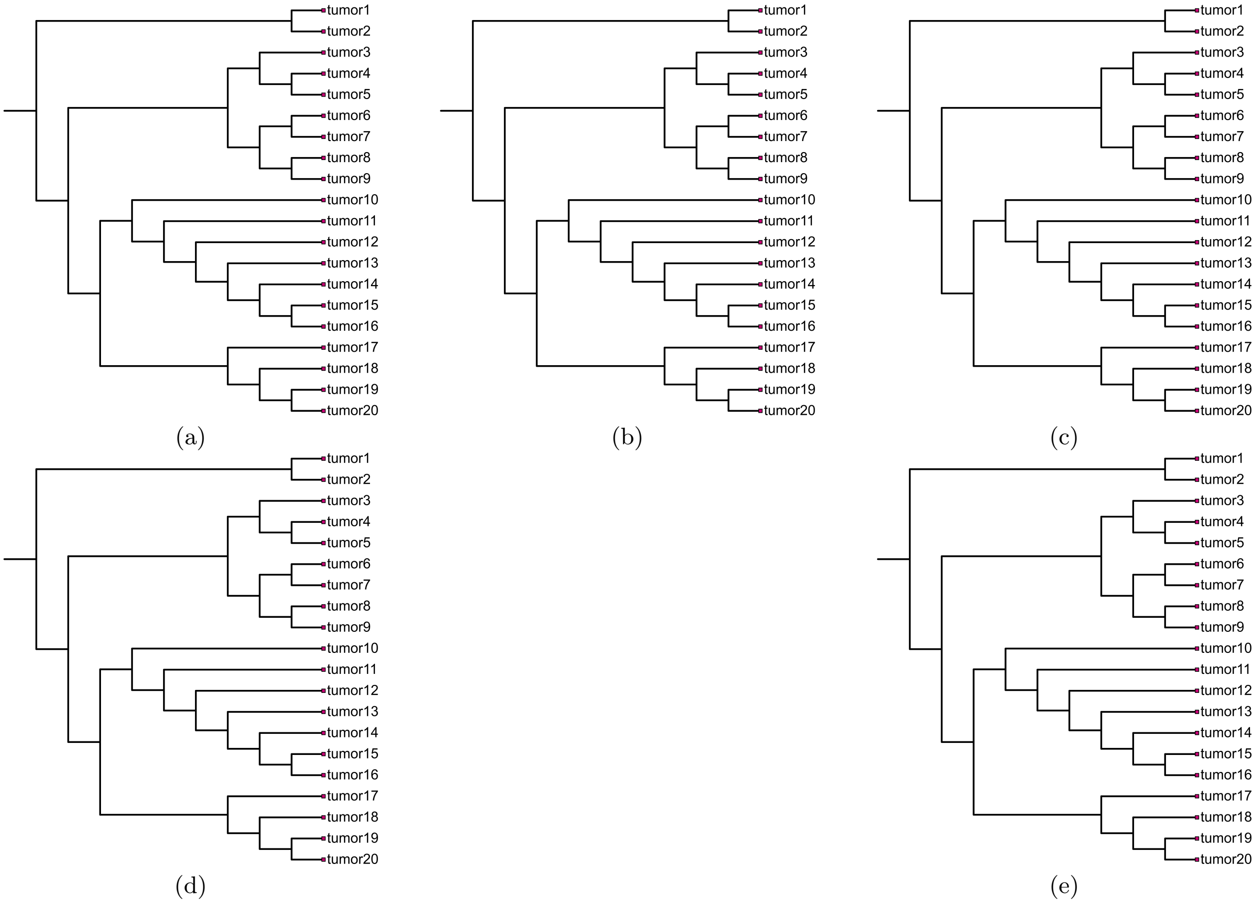


**Fig. S1. Phylogenetic trees constructed using simulated tumor genomes using GenCore.** Trees are constructed with (a) LCP level 4, (b) LCP level 5, (c) LCP level 6, (d) LCP level 7, (e) LCP level 8.

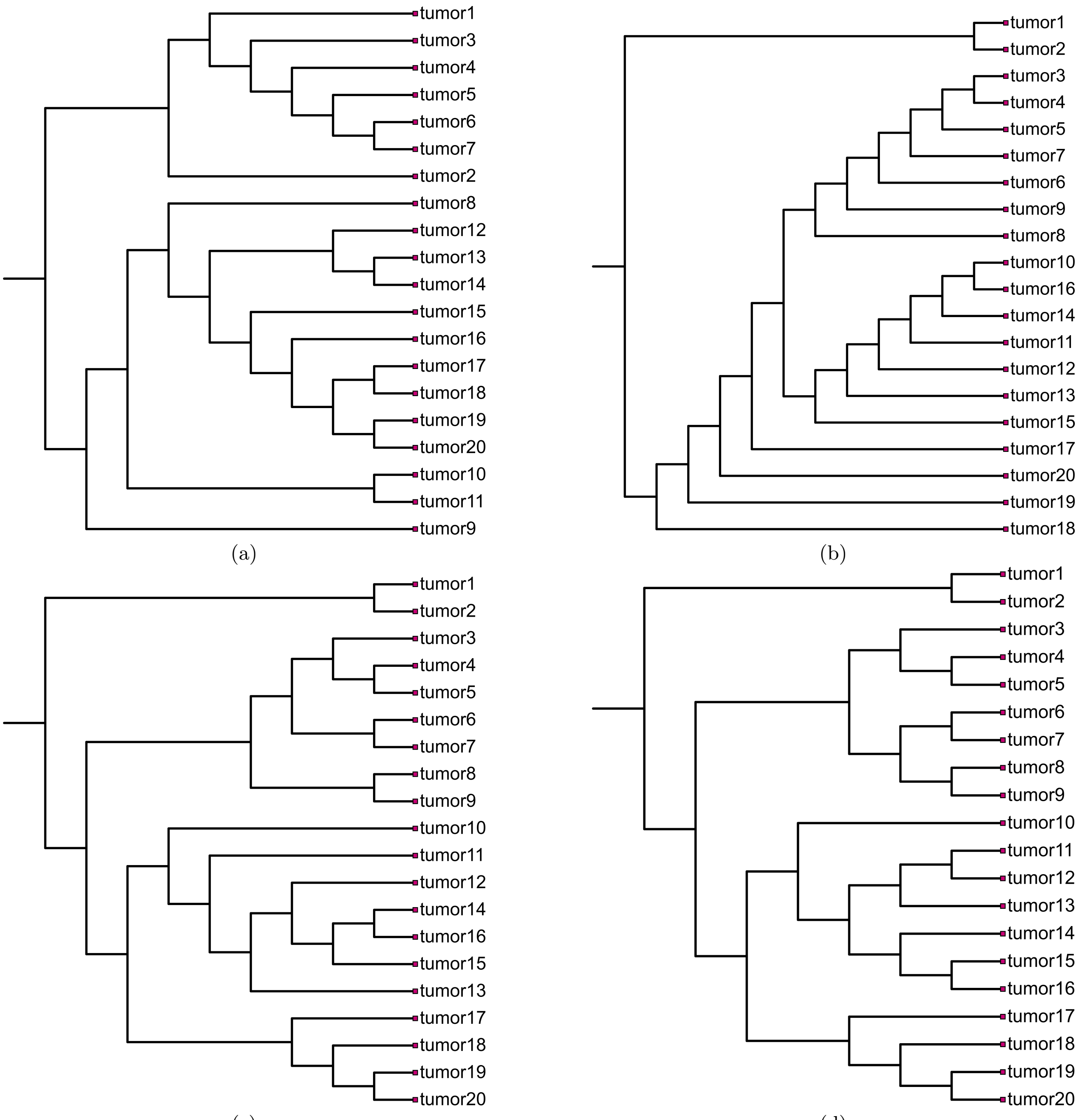


**Fig. S2. Phylogenetic trees constructed using simulated tumor genomes using `Mash` with k-mer size of 31.** (a) With sketch size 5,000; (b) With sketch size 50,000; (c) With sketch size 500,000; (d) With sketch size 5,000,000.

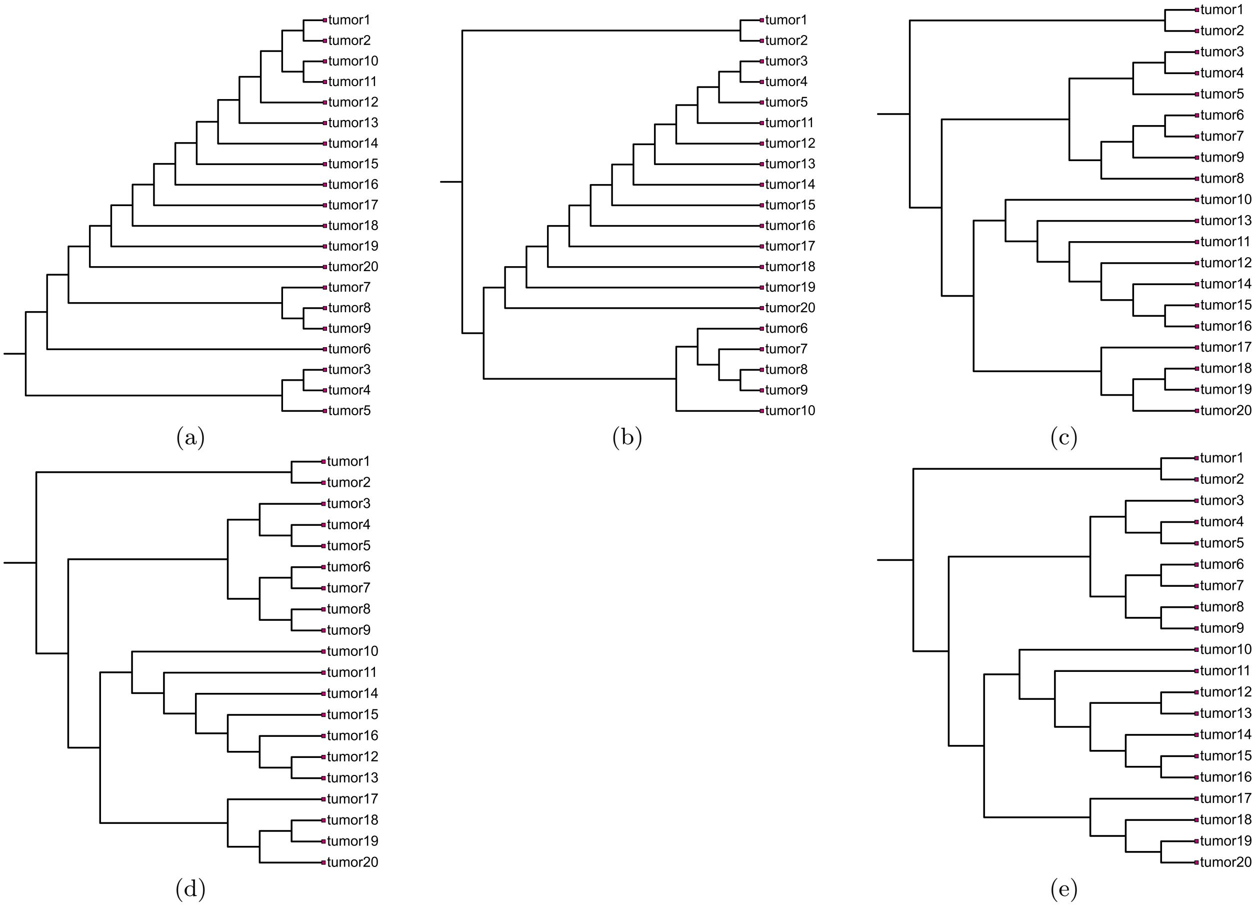


**Fig. S3. Phylogenetic trees constructed using simulated tumor genomes using `Dashing 2` with k-mer size of 21.** (a) With sketch size 1,000; (b) With sketch size 5,000; (c) With sketch size 50,000; (d) With sketch size 500,000; (e) With sketch size 5,000,000.

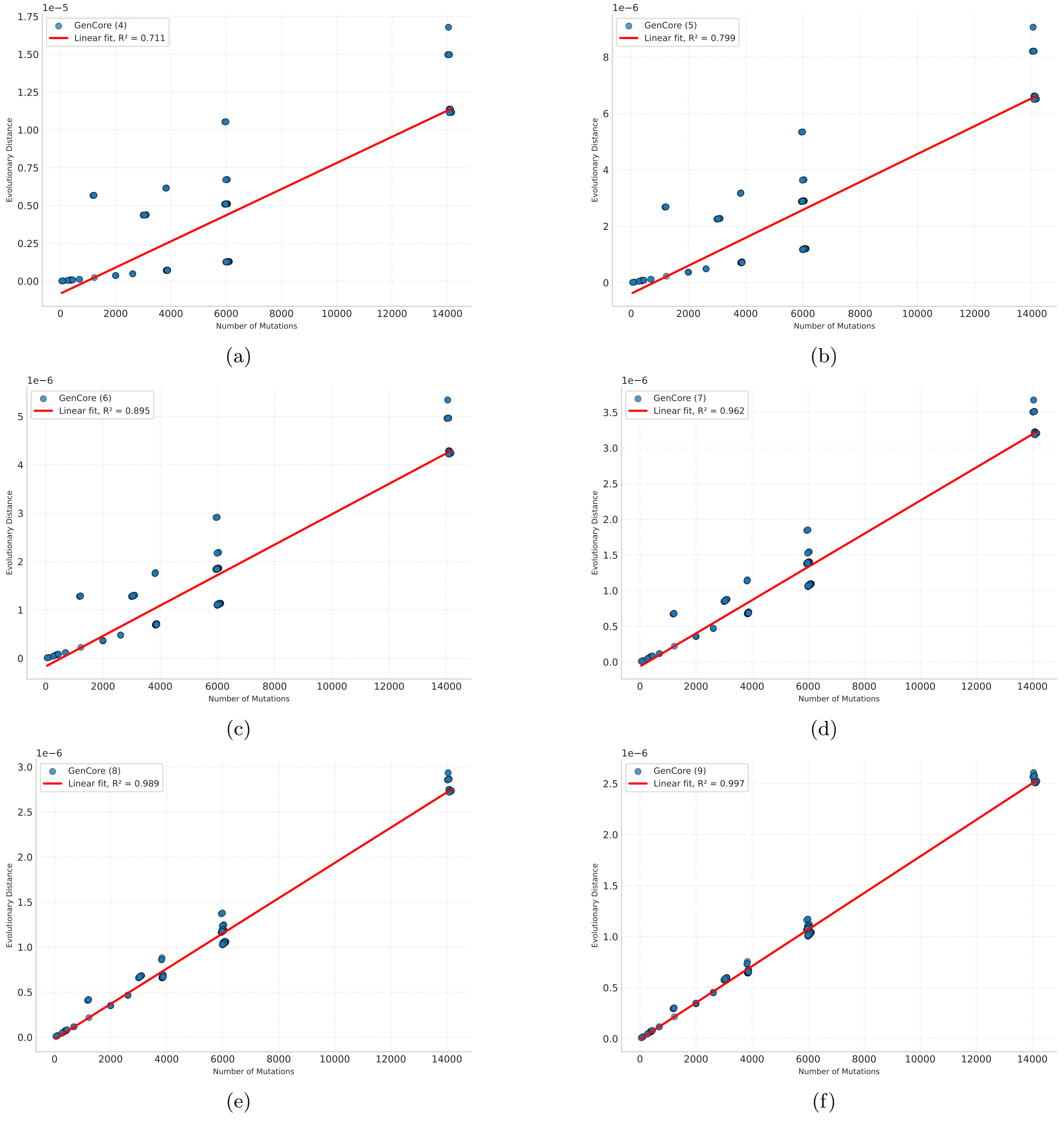


**Fig. S4.** ***p*-distance scores computed using GenCore vs number of mutations.** Plots are constructed with (a) LCP level 4, (b) LCP level 5, (c) LCP level 6, (d) LCP level 7, (e) LCP level 8, (f) LCP level 9.

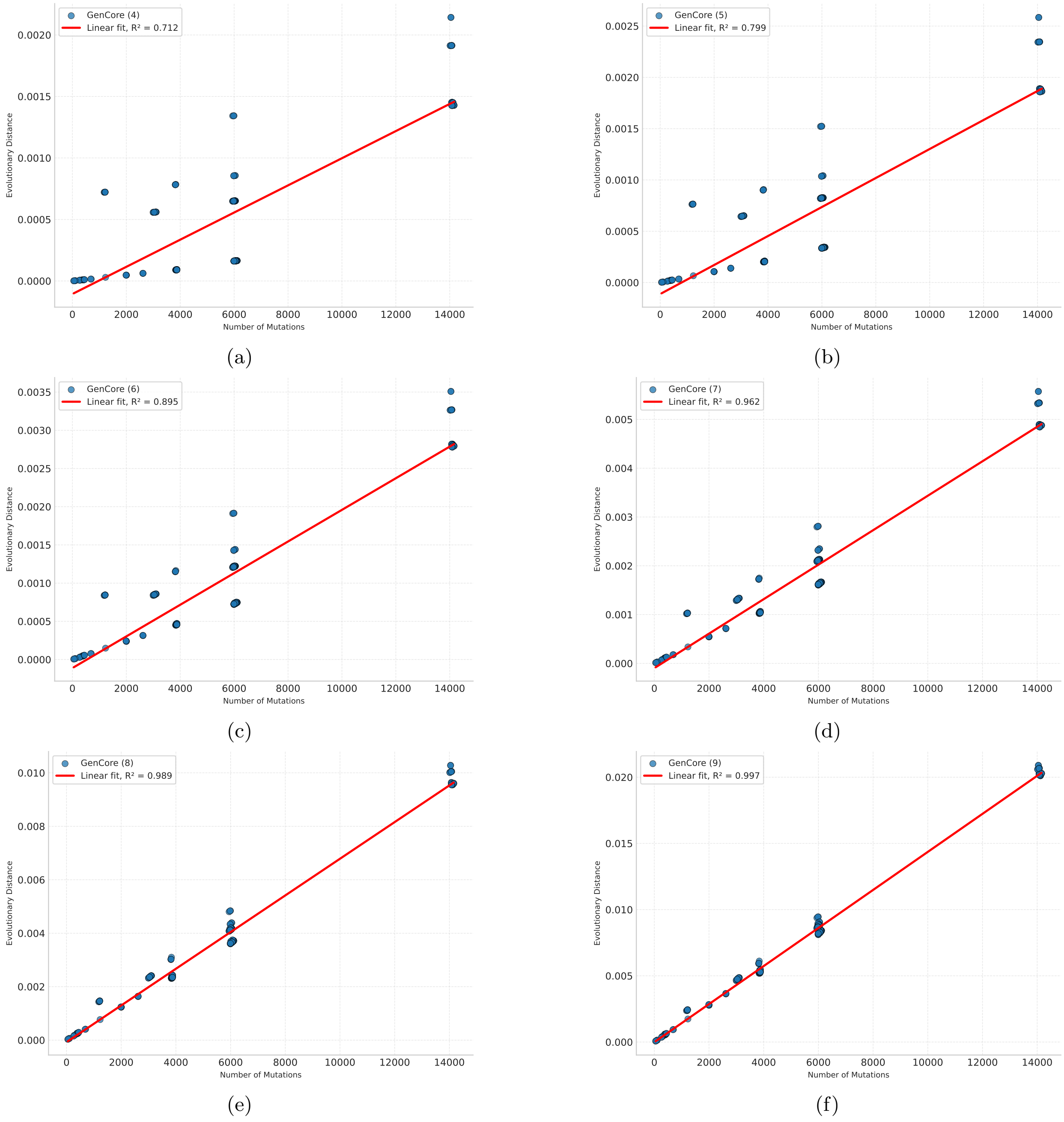


**Fig. S5. Jaccard-distance scores computed using GenCore vs number of mutations.** Plots are constructed with (a) LCP level 4, (b) LCP level 5, (c) LCP level 6, (d) LCP level 7, (e) LCP level 8, (f) LCP level 9.

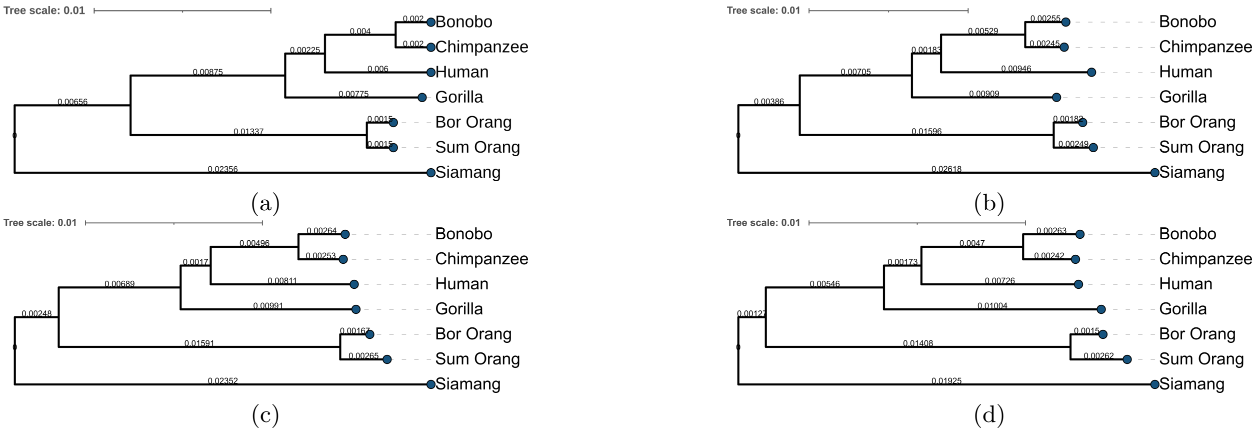


**Fig. S6. Pylogenetic trees constructed using complete primate genomes.** Panel (a) shows the known phylogeny, and Panels (b)–(d) show trees based on the distance matrix for levels 4-6.

**Table S1.** Summary of Primate Genomes

| Species | T2T Assembly Length (diploid) | Total Length HiFi Reads | Depth of Coverage |
|---|---|---|---|
| Gorilla | 6.91 Gbp | 375.6 Gbp | 54× |
| Bonobo | 6.29 Gbp | 199.2 Gbp | 32× |
| Chimpanzeee | 6.18 Gbp | 216.7 Gbp | 35× |
| Sumatran Orangutan | 6.26 Gbp | 317.0 Gbp | 51× |
| Bornean Orangutan | 6.23 Gbp | 193.0 Gbp | 31× |
| Siamang | 6.36 Gbp | 252.1 Gbp | 40× |
